# JAMS - A framework for the taxonomic and functional exploration of microbiological genomic data

**DOI:** 10.1101/2023.03.03.531026

**Authors:** John A. McCulloch, Jonathan H. Badger, Nikki Cannon, Richard R. Rodrigues, Michael Valencia, Jennifer J. Barb, Miriam R. Fernandes, Ascharya Balaji, Lisa Crowson, Colm O’hUigin, Amiran Dzutsev, Giorgio Trinchieri

**Affiliations:** Laboratory of Integrative Cancer Immunology, Cancer Immunobiology Section, Center for Cancer Research, National Cancer Institute, Bethesda, MD; Genetics and Microbiome Core, Laboratory of Integrative Cancer Immunology, Center for Cancer Research, National Cancer Institute, Bethesda, MD; Translational Biobehavioral and Health Disparities Branch, National Institutes of Health Clinical Center, Bethesda, MD

## Abstract

Shotgun microbiome sequencing analysis presents several challenges to accurately and consistently depict sample composition and functional potential. Here we present a two-part framework – JAMS (Just a Microbiology System) – whereby with raw fastq files and metadata as input, meaningful analysis within a sample and between a sample can be performed with ease for either shotgun or 16S sequences. JAMS is the first package to provide seamless deconvolution of functions into their taxonomic contributors. We validated our JAMS framework on two human gut shotgun metagenome test datasets against the popular tool MetaPhlAn 4. We further demonstrate the application of the JAMS package, particularly the plotting functions, on a mouse shotgun dataset.

## Introduction

High throughput shotgun sequencing has become an increasingly common and robust technique^1^ for examining microbial communities. The advantage of shotgun sequencing over 16S rRNA amplicon sequencing for profiling the microbiota in a sample is such that it gives not only a snapshot of microbial taxonomic diversity, but insights into functional potential ^2,3^ in diverse microbial populations such as those from the environment^4,5^, animal sources^6,7^, and humans ^8^. The development of second-generation sequencing which produces short, fragmented reads, presented yet another challenge for metagenomic analysis^9^. However, even with these advances in sequencing, much of the microbial world remains both taxonomically and functionally unknown^10–12^. Furthermore, our current understanding of microbes is heavily skewed, with 88% of microbial isolates belonging to only four bacteria phyla^11^. These are just some of the factors which need be considered when undertaking microbiome community analysis.

Shotgun sequencing profiling of the microbiome is more challenging, computationally and algorithmically, than 16S rRNA amplicon sequencing because unlike the latter, in which each 16S sequence type represents a single bacterial entity, shotgun reads may represent genomes incompletely, with swathes of reads coming from genomic regions which do not allow for accurate taxonomic classification of the organism they originated from. Different approaches have been developed to address this challenge. Reference-based approaches align reads to a representative reference^13^. However these approaches are time-consuming, leading to the development of clade-specific approaches which extract a unique subset of genes from the reference for classification at the read level ^14–18^. Methods such as these have apparent drawbacks when considering the vast majority of microbial members remain unknown^10–12^.

Another alternative is assembly-based approaches, whereby reads are assembled into longer sequences and then identified taxonomically^19^. The assembly-based approaches currently available, albeit few and far between, lag behind^20^. Furthermore, while there are a plethora of analysis options available, current approaches experience a performance drop particularly when taxonomically classifying at the species level^21^.

Moreover, the two main stages of microbiome project analysis are, namely, characterization of the microbiome profile within a sample, and thereafter comparison of microbiome structure across samples to find differences which may be relevant to the biology between sample groups. For shotgun sequencing analysis within a cohort or project, there is therefore a large number of steps which must be taken towards getting to data which allows for biological hypothesis testing and these steps are imbued with the burden of choice as to the exact tools and strategies implemented for achieving the objective. These choices are typically made by a single bioinformatician working on a single project, the consequence being that within a group or core shotgun analysis is liable to a much greater amplitude of heterogeneity than 16S sequencing which is usually obtained by using consolidated 16S sequencing analysis packages such as Qiime^22^, which has intra- and inter-sample analysis functionality as well as plotting capabilities.

In order to address this and other difficulties, we developed in house a software package named Just A Microbiology System (JAMS), which is an integrated framework of software applications for analyzing microbiology-related sequencing data in a user-friendly, consistent, reproducible and comprehensive manner. Staring with only raw fastq sequencing files and metadata, the JAMS package can, with minimal human interaction, yield meaningful and accurate results for shotgun sequences of either a microbiome or bacterial isolate samples. It can also process 16S amplicon sequencing data. JAMS also supports the importation of MetaPhlAn data into JAMSbeta. The JAMS package also includes highly customizable plotting and statistical functions for producing publication-quality heatmaps, ordination plots and bar graphs.

The JAMS package has been used for shotgun sequencing analysis since 2019 and the results it has yielded has been published in several peer reviewed papers ^6,23–30^. The code has been publicly available on Github since then at https://github.com/johnmcculloch/JAMS_BW. The repository is evolving and maintained with over 200 commits per year on average, and comprises currently of over 15000 lines of code.

The JAMS package contains a host of highly customizable plotting functions, which allow users to tie functionality back to taxonomy with ease. The JAMS package is unique in that it comprises a framework to easily mix and match the analysis of samples from different cohorts and projects including publicly available microbiome sequencing reads. It achieves this by dissociating the phases of within-sample analysis and between sample analysis. The two main scripts in the JAMS package (JAMSalpha and JAMSbeta) are dedicated to intra and inter-sample analysis, respectively. JAMSalpha yields a unique single file for each sample which can then be used by JAMSbeta to create plots and statistical analyses, thereby making it easy to add more samples subsequently without having to rerun the entire cohort in order to obtain feature-by-sample tables.

## Methods

### JAMSalpha

JAMSalpha is a single sample analyzer script for yielding .jams files (together with an informative pdf about the sample and quality/host filtered fastqs). The JAMSalpha command is executable from a UNIX command line and takes three argument groups. Namely, a sequence input; a path to a JAMS-compatible formatted database (see below); and arguments describing the nature of the sample, such as association to a host species or not, and if the sample is a metagenome, or bacterial isolate.

For the sequence input arguments, JAMSalpha can take short sequencing reads from the IonTorrent or any Illumina platform, the latter can be either paired end, mate-paired or single-ended reads. Alternatively, in place of specifying FASTQ read paths, it is possible to simply pass an NCBI SRA accession number, whose sequencing run will be downloaded and used as input. Assembled contigs or complete genomes in FASTA format can be used as input, as can a GenBank assembly accession number, in place of FASTQs with the consequent skipping of the read assembly step in JAMSalpha. In this case, the resulting analysis is of qualitative rather than quantitative value as feature coverage is not computed.

When running a JAMSalpha script with sequencing reads (or contigs) as input, it must be pointed to a pre-built JAMS-formatted database (JAMSdb), which we make available for download, containing a JAMS-compatible Kraken2 database for taxonomic classification, a curated taxonomy table and genome size information, Bowtie2 indices for *Homo sapiens* and *Mus musculus*, and accessory gene BLAST-based protein databases Resfinder^31^ and the Virulence Finder database -VFDB ^32^ made available through the Abricate GitHub repository ^33^.

Sequencing reads are quality-trimmed and adapter clipped using Trimmomatic^34^ and if applicable, then aligned to a host species genome using Bowtie2^35^ to eliminate host reads. Bowtie2 indices for *Homo sapiens* and *Mus musculus* are already included in the JAMS-compatible database and the user can just pass “human” or “mouse” or “none” to the -H argument of the JAMSalpha command. Alternatively, if a valid NCBI tax ID for a Metazoan organism is passed, JAMSalpha will download the single most complete most recent genome from that species, build a Bowtie2 index and use that to host-filter reads. Reads which are non-aligned to the host species (NAHS) are then used as input for assembly either by MEGAHIT^36^ for metagenomes or by SPAdes^37^ for bacterial isolates.

Contigs larger than 500bp are then classified taxonomically with Kraken2^38^ using a custom-built JAMS-compatible Kraken2 database. Although we plan to make available an updated Kraken2 database every 12 months, in order to reflect latest taxonomy and isolate references, there is included in the JAMS package is a script (JAMSbuildk2db) which runs on Unix command line and can be used for building customized JAMS-compatible Kraken2 databases. This script will download all genomes of Bacteria, Archaea, Fungi, Viruses and Protozoa excluding any genome with an entry on the “excluded_from_refseq” column on the https://ftp.ncbi.nlm.nih.gov/genomes/genbank/assembly_summary.txt manifest file, plus *Homo sapiens* genome GRCh38.p14 (NCBI accession number GCA_000001405.29) and *Mus musculus* genome GRCm39 (NCBI accession number GCA_000001635.9) and use these as input for building a custom Kraken2 database. We included the mouse and human genomes in the database in order to be able to classify and eliminate any contig resulting from the assembly of host reads which somehow survived bowtie alignment filtering to the host genomes.

Importantly, NCBI taxonomy is downloaded by the JAMSbuildk2db script and is “sanitized” by pruning taxonomic levels outside of Domain, Kingdom, Phylum, Class, Order, Family, Genus and Species. Strain level is included in the curated JAMS taxonomy table (JAMStaxtable) under a column named “IS1” for infra-species, but is not currently used. For species taxids which have a missing taxonomic level (as for instance viruses not containing phyla attributions), that taxonomic level for that taxid is annotatied with the word “missing”. JAMSalpha uses Kraken2 only to attribute NCBI taxids to sequences. The entire taxonomical lineage is obtained by looking up the taxid on the curated pruned JAMStaxtable. JAMS then computes the Last Known Taxon (LKT) which is the lowest non-missing, non-unclassified taxonomic level for a particular taxid. Because Kraken2 yields taxids based the highest weighted leaf of a taxonomic path, that is not always at the species level. Thus, some contigs may be terminally classified at a level higher than species, suggesting the presence of a microbial entity not present in the database, but whose lowest classified taxonomy can still be imputed by Kraken2 and if their contig sum length adds up to a higher proportion of the expected genome size for that taxon, the higher the confidence that the hit is not a mere false positive.

To enable this, the genome sizes of all genomes downloaded is calculated and the median genome size for each taxon at each taxonomic level is computed and included in the JAMS-formatted database. This is used by the JAMSalpha pipeline to estimate “genome completeness” of each LKT by dividing the contig sum length of all contigs classified as belonging to each LKT by the median genome size of that taxon. This is expressed as a proportion and values close to 1, or 100% suggest that the entire genome of the taxonomic entity is present and being evaluated. This concept is also used for filtration when analyzing differences between samples (see JAMSbeta section).

Furthermore, when building a new JAMS-compatible Kraken2 database using the JAMSbuildk2db script, it is possible to specify which organism groups and which assembly levels to include or exclude, if so wished, in order to build specific JAMS-compatible kraken2 databases.

We make available a full JAMS-formatted database, containing a 192 Gigabyte JAMS-formatted Kraken2 database, Bowtie2 indices for *Homo sapiens* and *Mus musculus*, plus BLAST-based accessory databases and the Silva v138 16S database which can be downloaded as a single tarball file at: https://hpc.nih.gov/~mccullochja/JAMSdb202212.tar.gz. This particular JAMS-compatible Kraken2 database contains 109003 unique NCBI taxonomy IDs (taxids), which are comprised of 71619 LKTs (a single LKT which is roughly equivalent to a species level classification may comprise more than one NCBI taxid). Of these, 39815 are bacteria, 25358 are viruses, 4523 are fungi, 1050 are archaea, 862 are protozoa and the few remaining are mouse or human or LKTs unclassified by name.

Contigs are annotated functionally using Prokka^38^. InterProScan^39^ annotation of the predicted proteome of each sample generated by Prodigal within Prokka is done if requested. InterProScan currently only runs on the JAMS_BW version of the JAMS package, on NIH’s HPC cluster, Biowulf as InterProScan in very computationally intensive. We are working on a solution to add InterProScan functionality outside of the Biowulf cluster, on any system which has InterProScan installed and running.

The reads which were used for assembly are then aligned back to contigs using Bowtie2 to gauge feature depth and the number of bases covering each contig is computed using Samtools^40^ and Bedtools^41^. The number of bases covering each individual annotated feature is also computed. JAMSalpha thus keeps tally of all bases covering an LKT by first identifying all contigs belonging to that LKT with Kraken2 and then summing all the bases covering these particular contigs. Unassembled reads can optionally be individually classified taxonomically using the same Kraken2 database and the number of bases for each NCBI taxid in unassembled reads can be added to the respective contig base coverage tally for that LKT. However, this option is deprecated as we have found that it does not increase the accuracy of LKT quantification especially if the overall read assembly rate is high (> 75%), so classifying unassembled reads is not default in JAMSalpha.

A flowchart describing the stages of the JAMSalpha pipeline can be seen in Figure 1.

**Figure 1:**
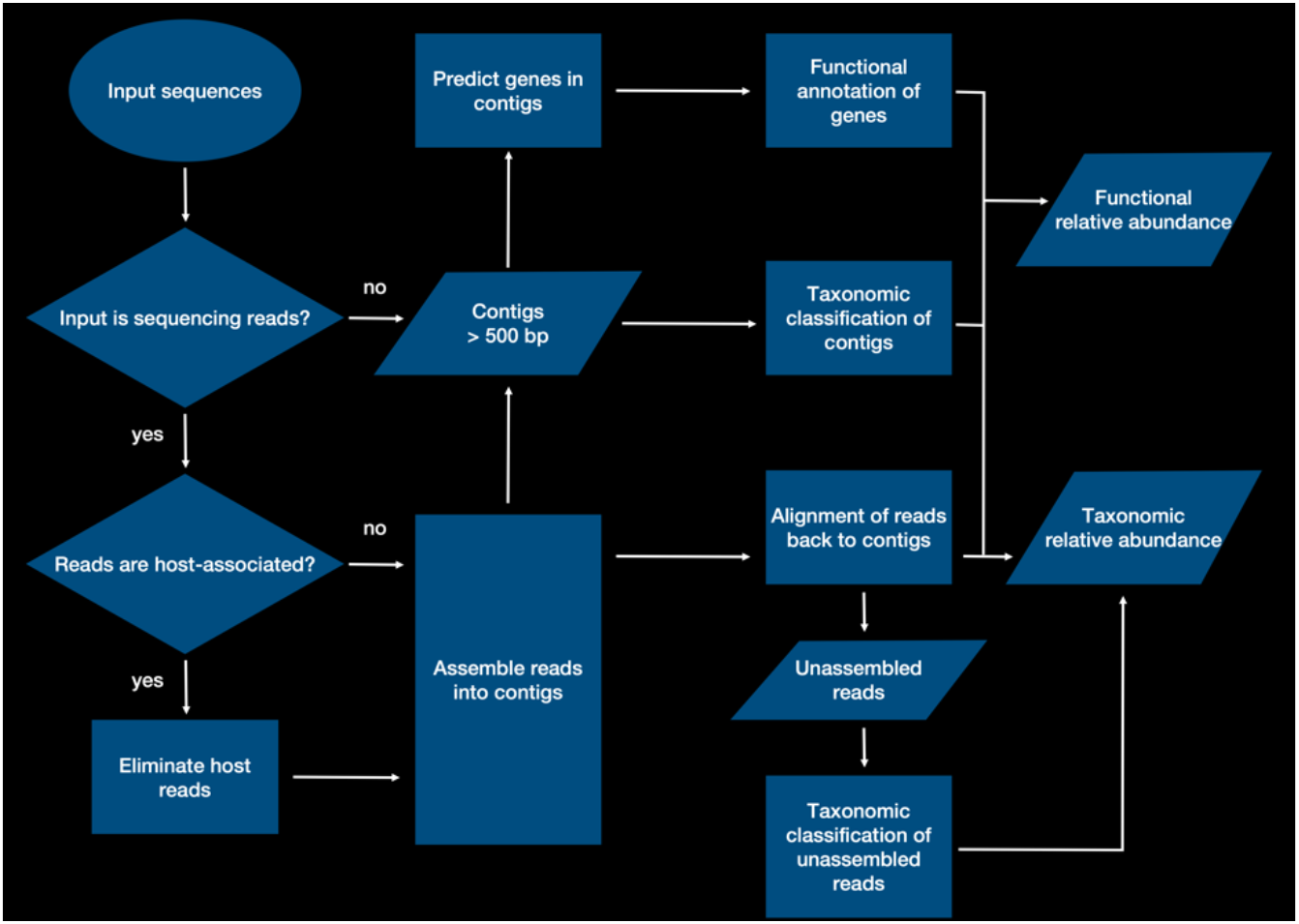
Flowchart describing each stage of the JAMSalpha pipeline.

### JAMSalpha output

At the end of a JAMSalpha run, host/quality filtered reads are saved to the output folder, a pdf document is generated containing JAMSalpha run analysis stats, sequencing read stats, contig assembly stats, an overview of the most abundant LKTs present in the sample in the form of a pareto chart, and tables with an overview of the functional characteristics of the sample. This report is usually more of use if the sample is an isolate and not a metagenome. Indeed, for isolates, a spreadsheet with all the functional and assembly characteristics of the sample is also generated. Most importantly, however, for all samples, a single file with the sample prefix and a .jams extension is generated. This file contains compressed R objects pertaining to all the quantitative and qualitative characteristics of sample, including tallies of base counts for each LKT found, as well as for each functional feature (gene) annotated, and the stratification by taxonomy of each functional annotation. This is possible because the accountancy of which feature (gene) came from which contig and what the taxonomy of each contig was. Also contained in a .jams file is information about read stats, assembly rates and estimated genome completeness of each LKT. However, a .jams file is not intended for human reading, rather, .jams files for individual samples can be compared to each other using the JAMSbeta script and metadata pertaining to the samples. See the JAMSbeta section for a detailed description.

### JAMSbeta

Comparison of samples with each other can be achieved by using the JAMSbeta script. This is an R script which can be executed directly on a Unix command line shell, and takes as input a folder path containing .jams files generated by JAMSalpha, and metadata in text format or in Microsoft Excel-compatible spreadsheet format (xlsx) mapping the prefix of each .jams file to metadata about the sample. JAMS-compatible metadata can really be any relevant information about the samples, either in discrete categories or in continuous variables as long as there is one variable in each of the metadata column plus a single column whose header is “Sample”, containing the .jams file prefixes. JAMSbeta will recursively look for matching .jams files in the folder path supplied and will extract the compressed R objects generated by JAMSalpha from these .jams files and use them to construct R objects containing counts tables.

After the JAMSbeta script has run, an R session image (.RData) is generated containing SummarizedExperiment objects^42^ each representing an analysis “space” of the samples. A SummarizedExperiment object is a unitary, subsettable object which ties together a features-by-samples table with annotation of features and annotation pertaining the samples. JAMSbeta constructs a single SummarizedExperiment object (SEobj) representing the qualitative and quantitative *taxonomic* properties of the samples with the following elements: a feature-by-sample assay matrix containing the number of basepair counts covering each LKT present in each sample; a feature-by-sample assay matrix containing the estimated genome completeness of each LKT in each sample computed by JAMSalpha as described above; a feature table containing the full taxonomic lineage of each LKT; the user-provided metadata for each sample; a class color dictionary specifying colors to be used for each discrete variable class of the metadata, in order to maintain colouring consistency throughout plots. This particular SEobj is named “LKT” because it represents only the taxonomic aspect of the samples.

In addition to the LKT SEobj, JAMSbeta also constructs an SEobj for gene product annotation obtained by Prokka in JAMSalpha for each sample. This is much like the LKT SEobj, except that features are gene product annotations rather than LKTs, and is thus a feature-by-sample assay matrix of basepairs covering all genes bearing each gene annotation. The associated feature table contains information about the gene product, such as gene symbol. The genome completeness assay is, of course, not present, but instead a feature-by-sample assay matrix containing the number of individual genes which bear each feature annotation within each sample. In addition to the class color dictionary, there is a sparse model matrix ^43^ stratifying the number of bases covered for each gene product annotation by LKT. In other words, with the “Product” SEobj, it is possible for downstream JAMS plotting functions to split a particular functional gene annotation into their relative contributing taxa and, thus, not only find that a gene product is enriched in a group of samples, but also which taxa bear genes annotated with this particular gene product within each sample.

SEobj generation does not stop at LKT and Product, analogous SummarizedExperiment objects are generated also for Enzyme Commission numbers (ECNumbers) and for the BLAST-based protein databases present in the JAMS database, such as antibiotic resistance genes from the Resfinder database and virulence genes from the aforementioned VFDB database. If InterProScan is run on the JAMSalpha phase for the samples, then SummarizedExperiment objects representing InterProScan analysis spaces are also generated, also with stratification of InterPro signatures into their contributing taxa.

All these SummarizedExperiment objects can be used as input for one of six JAMS plotting functions to generate publication-quality plots of any aspect of the microbiome of the samples being analyzed.

*plot_Ordination* will generate ordination plots using either PCA, PCoA, tSNE or UMAP as dimensionality reduction algorithms. Several options regarding how the ordination plot sample dots are to be colored, shaped, joined, have centroids drawn up, among other options, are easily implementable by just naming the relevant metadata columns.

*plot_relabund_heatmap* will generate heatmaps of features by samples based on the ComplexHeatmap package^44^. A panoply of different heatmap styles can be generated easily by just passing to the function arguments which metadata column names are to be used for heatmap annotation, which one is to be used for statistical significance testing using the Mann-Whitney-Wilcoxon test or ANOVA, with heatmap colors either by relative abundance in Parts per Million or scaled Z-score, plus a host of other aesthetic options. Log2 fold changes and raw and adjusted p-values are automatically calculated and added to the heatmaps. Naïve clusterization or heatmap group splitting can be imposed.

*plot_relabund_features* will generate boxplots of features by sample groups. Stats of comparisons between groups using the Mann-Whitney-Wilcoxon test or ANOVA are calculated internally and raw and adjusted p-values coupled with log2 foldchange values when applicable are drawn up on the plot for comparing a single taxonomic or functional feature between groups or aggregated features. Furthermore, when plotting a functional space such as Product, ECNumber or any InterPro signature, stratification of the functional feature into the contributing taxa in a second boxplot is easily achievable with a single argument within this plotting function.

### Benchmarking JAMSalpha

#### *Mock Com*munity Samples *used for assessment*

The two mock communities used for the analysis in this report were previously published ^45,46^ and available for download at the Sequence Read Archive at NCBI. The mock microbiome community used by Tourlousse et. al., represented a near even concentration of 20 bacterial species and 20 genera prevalent in the human gut microbiome (Table 1A). The 6 paired-end fastq were available at NCBI SRA under BioProject PRJNA747117 (SRA accession numbers: SRR17380241-SRR17380246). The analysis in this report will focus on 19 species as the species and the strain of *Bifidobacterium longum* and *Bifidobacterium longum subsp. longum* were collated into only one species.

**Table 1A:** Expected Species and relative abundances of Tourlousse mock community

| Species Name | Expected Relative Abundance | Taxonomic ID |
| --- | --- | --- |
| *Akkermansia muciniphila* | 6% | 239935 |
| *Anaerostipes caccae* | 5% | 105841 |
| *Bacillus subtilis* | 5% | 1423 |
| *Bacteroides uniformis* | 5% | 820 |
| *Bifidobacterium longum* | 10% | 216816 |
| *Blautia sp.* | 5% | 2877527 |
| *Collinsella aerofaciens* | 6% | 74426 |
| *Cutibacterium acnes* | 5% | 1747 |
| *Enterocloster clostridioformis* | 5% | 1531 |
| *Escherichia coli* | 6% | 562 |
| *Flavonifractor plautii* | 4% | 292800 |
| *Lactobacillus delbrueckii* | 4% | 1584 |
| *Megamonas funiformis* | 4% | 437897 |
| *Megasphaera massiliensis* | 5% | 1232428 |
| *Parabacteroides distasonis* | 5% | 823 |
| *Pseudomonas putida* | 4% | 303 |
| *Ruminococcus gnavus* | 6% | 33038 |
| *Staphylococcus epidermidis* | 5% | 1282 |
| *Streptococcus mutans* | 7% | 1309 |

**Table 1B:**
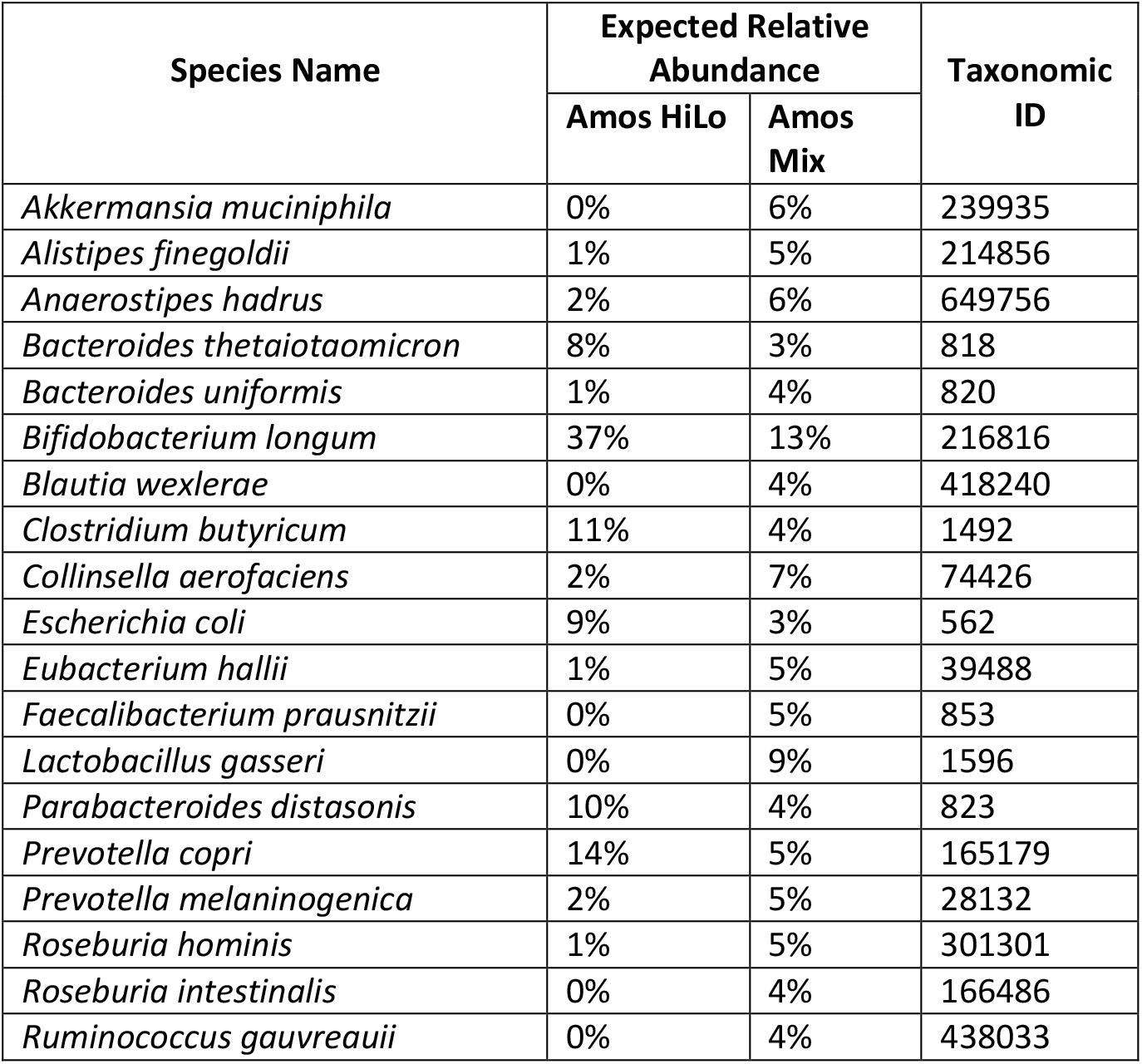
Expected Species and relative abundances of Amos mock communities

The second set of mock communities included in this report, previously published by Amos et. al, were 5 even (Amos Mixed) and 5 staggered (Amos HiLo) mixtures of representative gut microbiome species (Table 1B). These communities contained 19 species and 16 genera. The 10 paired end fastq files were directly downloaded under BioProject ID PRJNAA622674 (SRA accession numbers: SRR11487931-SRR11487935 and SRR1148793-SRR11487941).

### Quality control checking of mock community samples

All mock community samples were submitted to quality control checking using FastQC ^33^ and trimming procedures were carried out using fastp (https://github.com/OpenGene/fastp). FastQC was run using the default parameters in version 0.11.9. For the Tourlousse dataset, the following fastp parameters were used: -q 15, --cut_front, --cut_tail. The Amos HiLo and Amos Mixed samples were trimmed using the following fastp parameters: -q 25, --cut_front, -- cut_tail, -l 100, -f 15 and -t 75.

### Biobakery Workflows Pipeline for Shotgun Metagenomics processing of mock samples

The metagenomic assembly workflow used in this report for comparison to the JAMS package was based on the updated version of Biobakery Workflows which includes MetaPhlAn 4^14,48^. Briefly, this workflow was installed on the NIH High Performance Computing environment Biowulf (https://hpc.nih.gov/systems/), which utilizes a slurm workload manager. The installed Biobakery workflows package, downloaded and installed directly (https://github.com/Biobakery/Biobakery/wiki/Biobakery_workflows), was used in part to obtain the taxonomic profiles for the analysis. The tools utilized from the Biobakery workflows package were the kneaddata tool, used for decontamination, trimming, and quality control, and MetaPhlAn4 for taxonomic profiling. MetaPhlAn4 is based on a curated database of 1.01 million prokaryotic reference and metagenome-assembled genomes. The MetaPhlAn4 profiler fundamentally differs from the kraken2 classifier used by the JAMS package as MetaPhlAn4 is based on a marker gene analysis. For more details, refer to the publication by Segata, et. al. The command used to obtain the taxonomic profile for the mock communities was as follows: Biobakery_workflows with the wmgx flag. The results from using this package will be referred to as ‘Biobakery4’ in this report.

After running the workflow in Biobakery workflows, the metaphlan_taxonomic_profiles.tsv from the merged metaphlan directory in the Biobakery workflows output was cleaned into the species_relab.txt file using a grep command. Separate tables for the genus level and the species level features were created, where $1 is the merged table and $2 is the output path:

Genus:

~~~
grep -E ”(g_)|(taxonomy)” ”$1” > ”$2”
~~~

Species:

~~~
grep -E ”(s_)|(taxonomy)” ”$1” > ”$2”
~~~

Both genus and species-level relative abundance tables were parsed using a custom python script. The customized python script cleaned the table and attached TAXIDs from the NCBI Taxonomy database (https://pubmed.ncbi.nlm.nih.gov/22139910/). This yielded the final relative abundance tables used for the analysis in this report.

### JAMS Processing for Shotgun Metagenomics processing of mock samples

The alignment database was generated as previously described and for the purpose of the mock community processing included in this report, the updated December 2022 database, deemed ‘JAMSdb202212’ version is used for assessment in this report (available for download at https://hpc.nih.gov/~mccullochja/JAMSdb202212.tar.gz).

The commands used for the JAMS processing workflow were as follows:

First, to run JAMSalpha:

1)The JAMSmakeswarm utility was used to generate the swarmfile for the job submission:

~~~
JAMSmakeswarm -r [path/to/reads] -d [path/to/db]
~~~

Which created a swarmfile where each line contained the following JAMSalpha command for each sample:

2)Each sample was submitted using the following command in a swarmfile.

~~~
JAMSalpha -f [path/to/f_read] -r [path/to/r_read] -o
[path/to/output] -H human -p [prefix]
~~~

3) The swarmfile was submitted:

~~~
swarm -g 246 -t 56 --time=24:00:00 --module R,samtools --
gres=lscratch:400 -f JAMS.swarm
~~~

Once JAMSalpha completed, JAMSbeta was run to generate the relative abundance tables using the following command:

~~~
JAMSbeta -p [projectname] -o [output] -t [metadata].tsv -y
[path/to/jamsfiles] -e -k -z -n
LKT,ECNumber,Product,Pfam,Interpro,GO,resfinder
~~~

For JAMS, the excel file from JAMSbeta was parsed using a custom script. It searched for rows tagged with s_ (species level) or both s_ and g_ (for genus level), and also the “Unclassified” row. Both genus and species-level relative abundance tables were parsed using the custom python script which attached TAXIDs from the NCBI Taxonomy database (https://pubmed.ncbi.nlm.nih.gov/22139910/). In order to obtain the genera names from species, the species scientific names were split and the first part (i.e., the genus) was used. These feature tables were then processed in the JMP Statistical Computing Software version 16 (SAS Headquarters, Cary, NC).

### Assessing Accuracy between the observed and expected communities

The Aitchison Distance (AD) is a measure of the Euclidean distance between centralized log ratio (CLR) transformed values and has been shown to be a superior metric for assessing distance between compositional data^49^. The AD was calculated between the observed and expected relative abundance values in several steps. Since some expected features were missed by the two pipelines, and some pipelines reported features that were not expected (false positives), the outer join created a matrix with some missing values. Missing values were replaced with zeroes, a zero-replacement strategy was needed in order to perform the CLR transform. The following strategy was followed to replace zeros in the full data matrix. For both the expected and the observed the minimum non-zero value was determined. A uniform draw between 10 percent of the minimum and the minimum value was used for zero replacement of both the expected and the observed^50^:

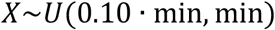

All other metrics were calculated without replacement of zeroes (on the non-CLR transformed data). Since the uniform draw method is a stochastic process, the statistics were calculated 5 times for each sample. Since each sample is also a replicate, all of the runs were averaged together for the final AD.

For this analysis, all relative abundance values less than 0.01% were removed before assessing closeness between the observed and expected values. The relative abundances were then renormalized to 100%. Expected and observed taxa were merged using TAXID in order to assess how well each pipeline characterized the mock communities at both the genus and species level, which was performed to control for divergences in naming. Relative abundance plots were created for all mock communities included in the report. The total False Positive Relative Abundance (FPRA) rate was computed by taking the total sum of the relative abundance of the false positives from the outer joined data matrix [add citation from Amos paper, see first comment]:

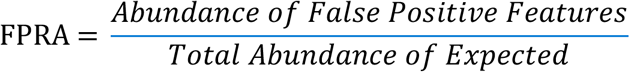

The reported sensitivity (Sens) metric in this report indicates the number of observed genus or species that are expected (i.e., true positives) over the total number of expected genus or species. All replicate mock communities were averaged in order to display one result for the three included communities.

The Sensitivity (Sens) was computed as the number of species correctly identified divided by the total number of expected species. Rows were selected where **both** the expected and observed RA were greater than zero (corresponding to a hit).

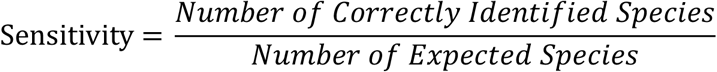

### JAMSbeta example

We generated Illumina shotgun sequencing data (deposited in SRA under BioProject accession number PRJNAXXXXXXX) from a simple SPF mouse experiment where animals were separated into two groups of five individuals. One group was treated orally with vancomycin and the other group was treated with water. We used JAMS to determine the differences in the microbiota after antibiotics were given. Samples were taken at two timepoints: day 1 before antibiotics and 5 days after antibiotics.

To run JAMSalpha, the following commands were run:

1)The JAMSmakeswarm utility was used to generate the swarmfile for the job submission:

~~~
JAMSmakeswarm -r [path/to/reads] -d [path/to/db] -H mouse
~~~

Which created a swarmfile where each line contained the following JAMSalpha command for each sample:

~~~
JAMSalpha -f [path/to/f_read] -r [path/to/r_read] -o
[path/to/output] -H mouse -p [prefix]
~~~

2) The created swarmfile was submitted:

~~~
swarm -g 246 -t 56 --time=24:00:00 --module R,samtools --
gres=lscratch:400 -f JAMS.swarm
~~~

After the successful running of JAMSalpha and the creation of .jams files, JAMSbeta was launched with the following command:

~~~
JAMSbeta -p [projectname] -o [output] -t [metadata].xlsx -y
[path/to/jamsfiles] -n
LKT,ECNumber,Product,Pfam,Interpro,GO,resfinder
~~~

JAMS includes several functions which are integrated into the plotting functions to achieve elimination of spurious taxonomic features, or to filter above a threshold. Relative abundance is expressed in Parts per Million (PPM) which is obtained for each feature, be it taxonomic or functional, by normalizing the number of bases for a feature in a sample by the total number of bases sequenced in that sample and multiplying by a factor of 10^6^. This is done every time a JAMS plotting function is called, as the SEobj contain raw base counts and not relative abundances. The total sequencing effort in base counts for each sample is contained within every JAMS SEobj. Feature filtering is achieved by specifying a minimum relative abundance PPM threshold in at least x % of samples for a feature to be kept after filtering. Thus, the “*featcutoff =*“ argument in JAMS plotting functions takes two values, namely, the PPM threshold and the percentage of samples that have to cross that threshold. So featcutoff *=c(50, 15)* means discard any feature that does not occur in at least 50 PPM in at least 15% of samples in the SEobj being passed to the plotting function. For the taxonomic LKT SEobj, the “*GenomeCompletenessCutoff =*“ also takes a double value, the first being for minimum percent estimated genome completeness threshold and the second being the percentage of samples which have to cross that threshold.

Titration of filtration thresholds show that, in a cohort of 39 mouse fecal samples used as example data the number of LKTs drop precipitously to relatively stable levels very fast (Figure 2). When paired with genome completeness filtration, the effect is even starker, with rapid stabilization of the number of surviving features even with very small PPM filtration values (Figure3)

**Figure 2:**
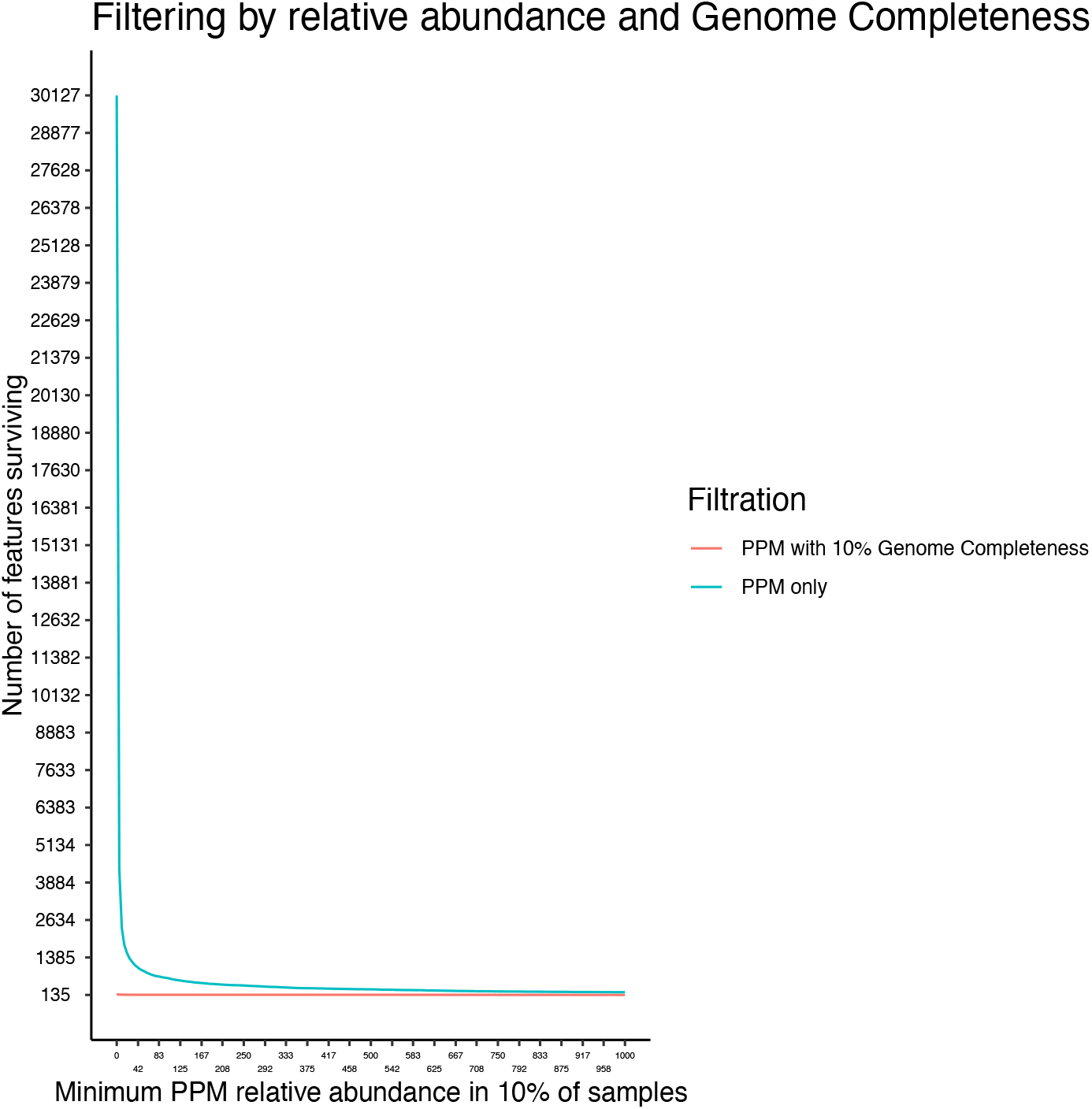
Number of surviving LKTs with increasing filtration of PPM relative abundance thresholds in at least 10% of samples.

**Figure 3:**
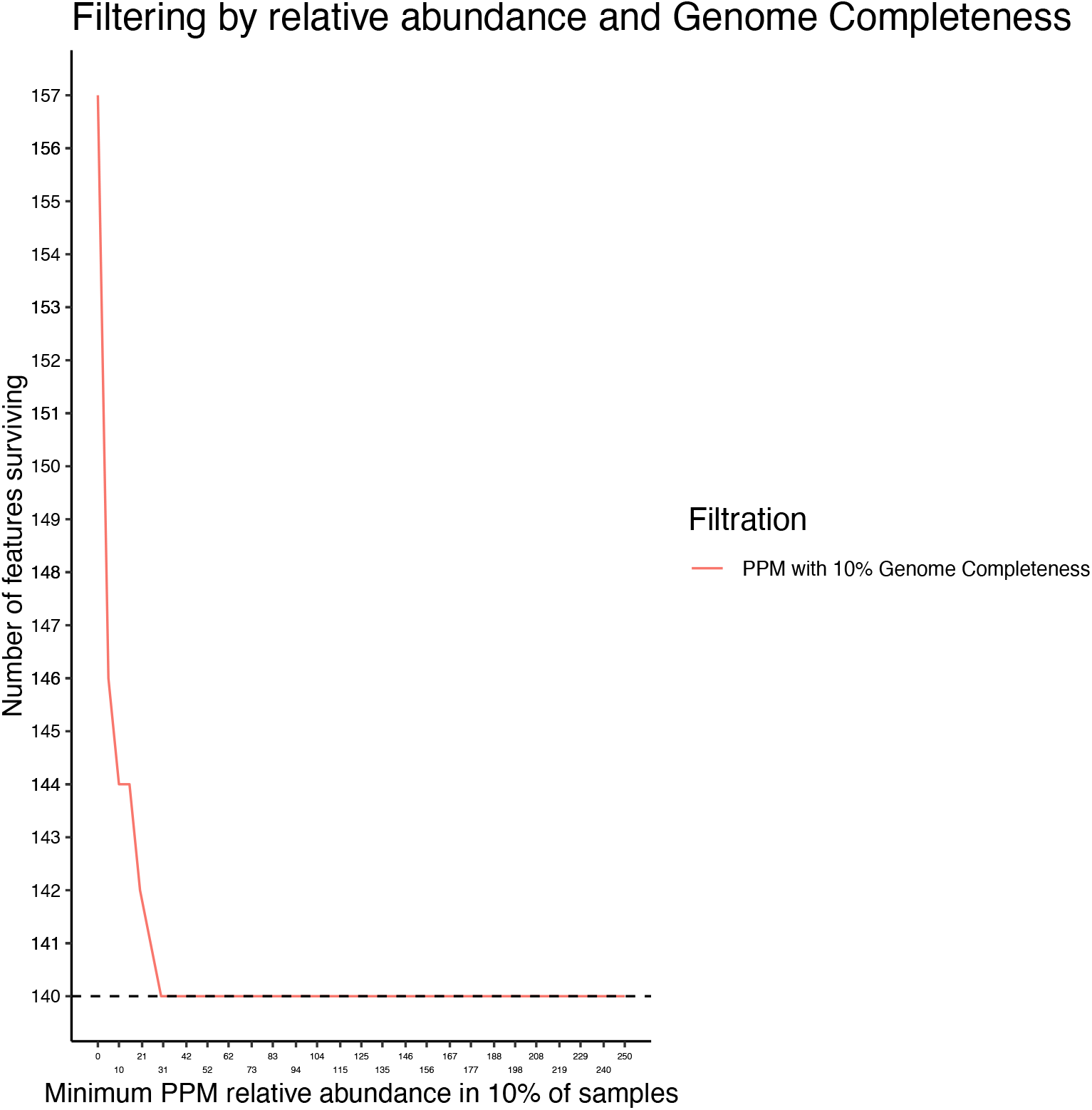
Number of surviving LKTs with increasing filtration of PPM relative abundance thresholds in at least 10% of samples whilst maintaining a fixed genome completeness threshold of at least 10% genome completeness in 10% of samples.

**Figure 4:**
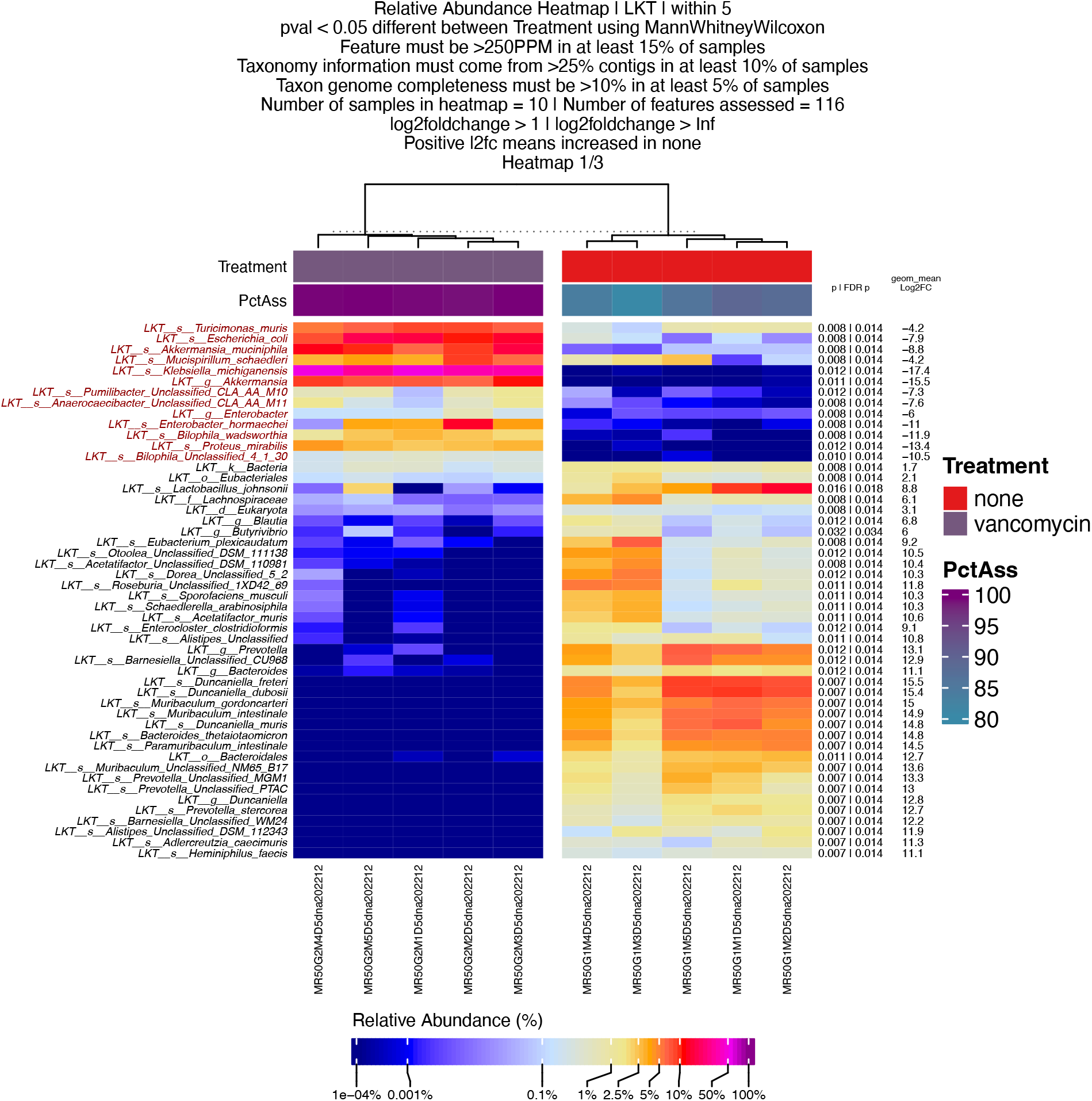
Example relative abundance heatmap of stool samples of mice treated (left) or not (right) with oral vancomycin. This plot was generated with a single line of code from a JAMSbeta session. P-values are obtained with Mann-Whitney-Wilcoxon U-Test and p-value adjustment was obtained by FDR. The log2foldchanges were calculated with the log of the foldchanges between the geometric mean relative abundance of each feature within each group.

**Figure 5:**
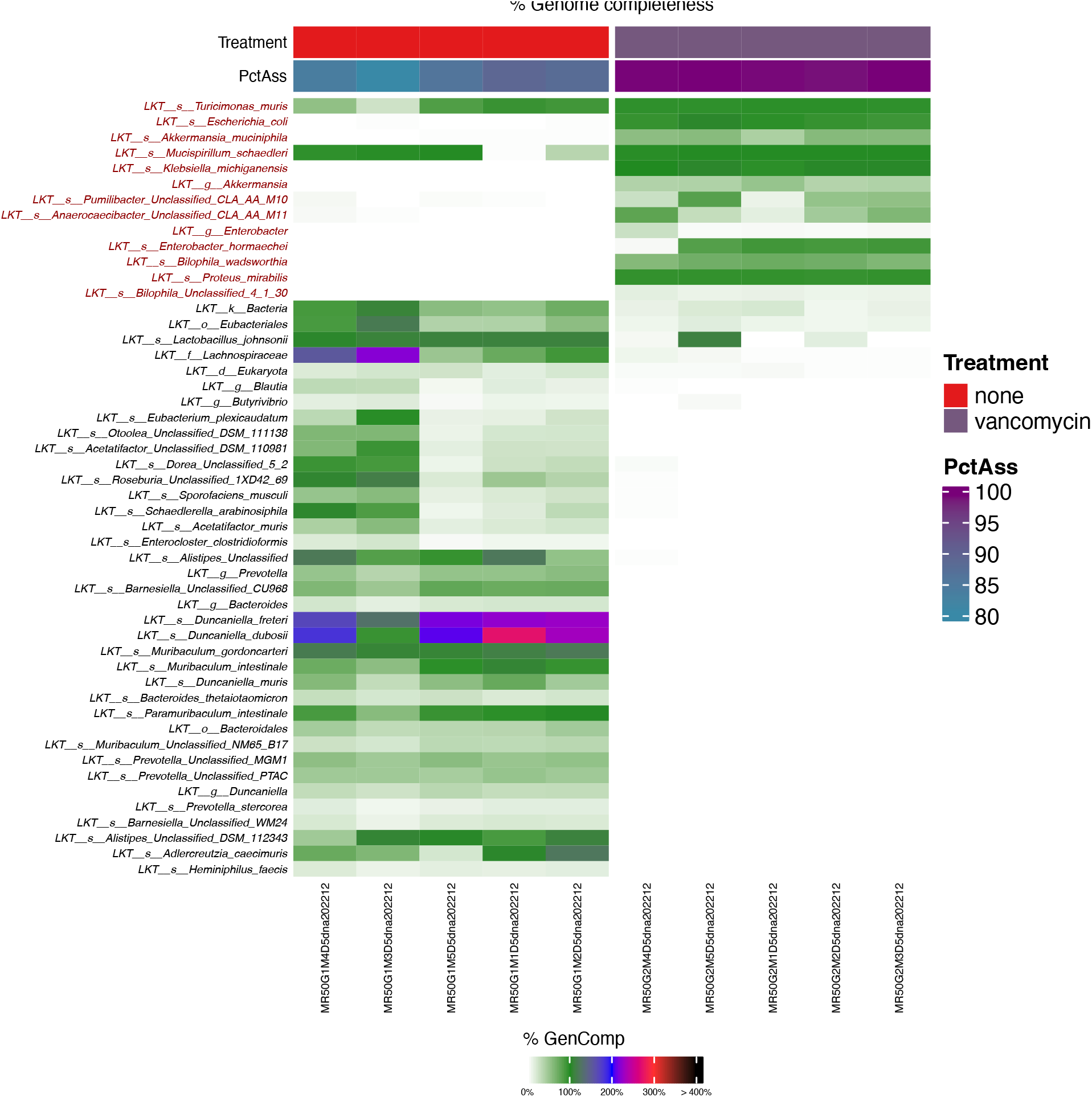
Example relative abundance heatmap of stool samples of mice treated (left) or not (right) with oral vancomycin. The feature and sample order are the same as in Figure 4, but cells represent the estimated Genome Completeness of each LKT within each sample.

**Figure 6:**
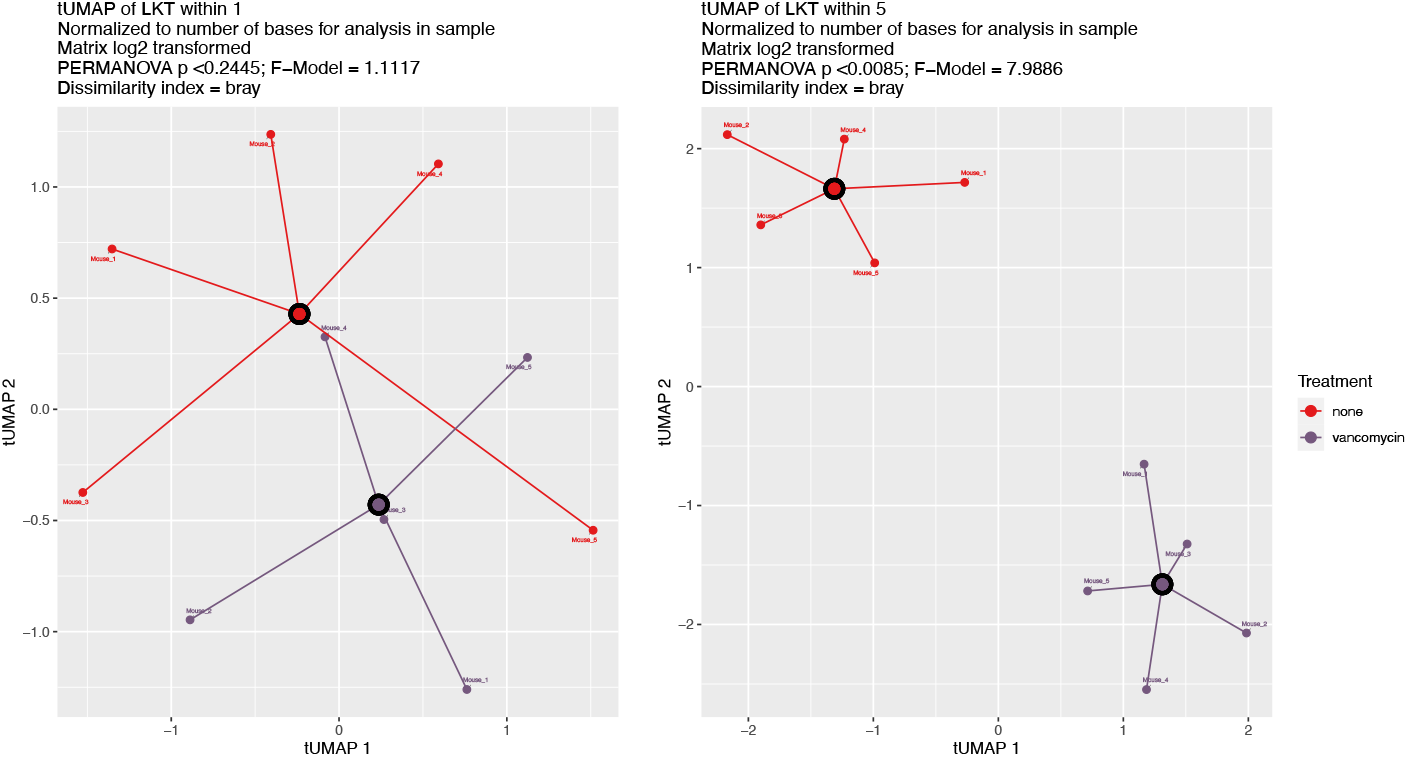
Example ordination plot (tUMAP) of LKT in stool samples of mice treated (purple) or not (red) with oral vancomycin with centroids plotted as well. On the left, is a tUMAP of the LKTs within the samples before starting treatment on day 1. On the right, is a tUMAP of the LKTs within the samples after 5 days of treatment. This plot was generated with a single line of code from a JAMSbeta session.

**Figure 7:**
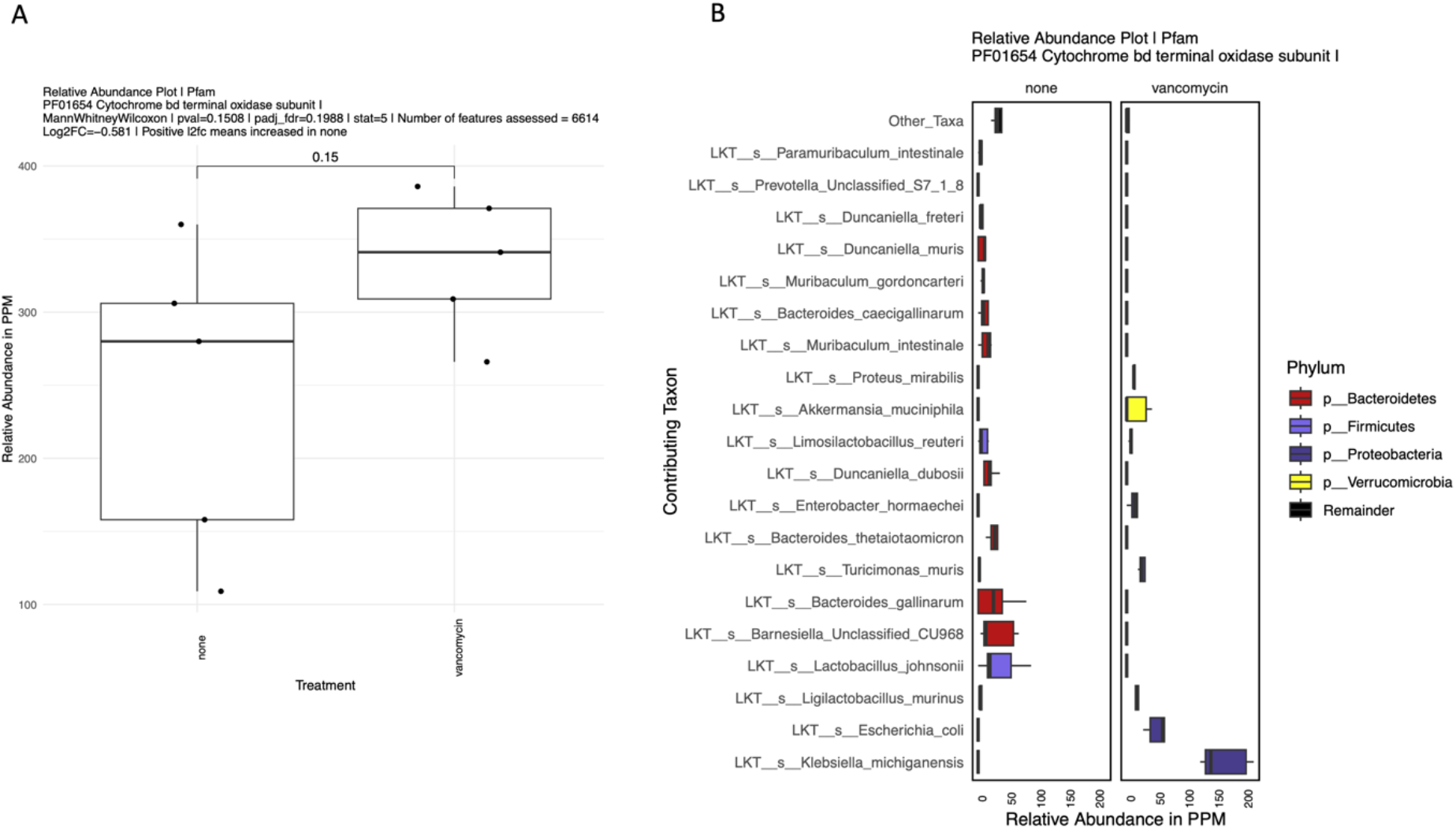
Example relative abundance feature plot comparing relative abundances of cytochrome bd terminal oxidase subunit I in mice treated or not with oral vancomycin after 5 days of treatment. Panel A is a box plot of the overall relative abundance of this feature in samples. Panel B are boxplots representing the taxonomic stratification of the plot in panel A, depicting the relative abundances of cytochrome bd terminal oxidase subunit I within each contributing LKT which bears this feature.

## Results

### Mock Community Read Statistics

The sequencing platform used for all mock communities was NextSeq 500 (Illumina, USA) (Table 2). The average total number of sequences in the Tourlousse data set was 5.9M, average read length was 145 basepairs (Table 2). The average total number of sequences for the Amos HiLo mock community was 1.9M and the average read length was 200 basepairs. The average total number of sequences for the Amos Mixed mock community was 2M with average read length of 200 basepairs.

**Table 2:**
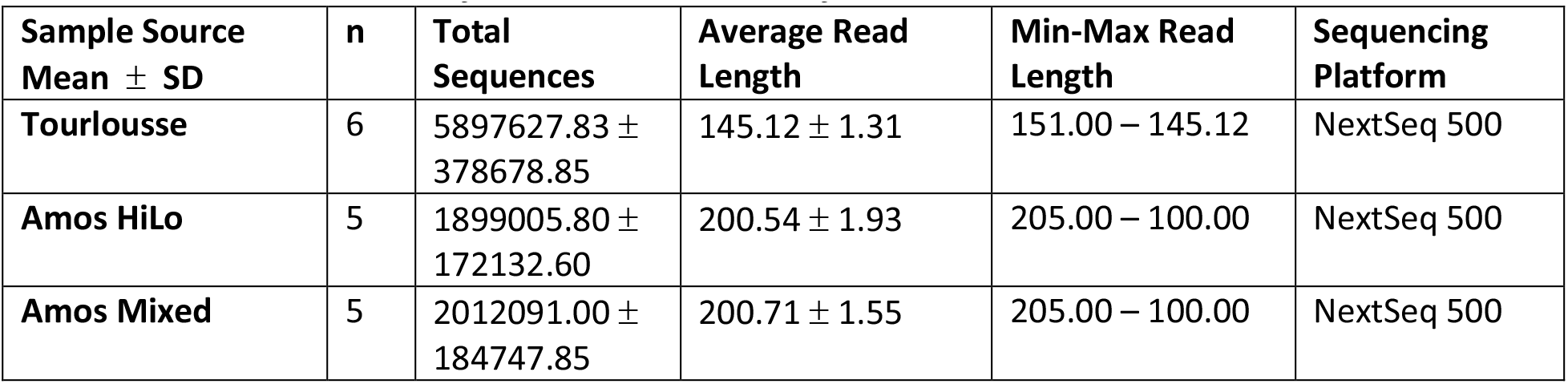
Raw Read Statistics for the Communities of Interest

### Performance of Metaphlan and JAMS package

The observed relatives abundance values were compared to the expected relative abundance values for each mock sample from the two packages (Figures 8-10). For the Tourlousse samples, Biobakery4 was able to find 18/19 of the genera (missing *Ruminoccoccus*), whereas JAMS were able to find all 19 genera in the mock community samples (Figure 8A). At the species level, Biobakery4 was unable to detect *Blautia sp* NBRC-113351 while JAMS showed very low abundance of this species (0.002%) (Figure 8B). Biobakery4 found species *Blautia producta* at .047% relative abundance which may have been misclassified as the wrong species. Biobakery4 had a total of 4.72 ± 0.04 FPRA at the species level while JAMS had a total of 7.45 ± 0.22 FPRA at the species level (Table 3B). Biobakery4 showed an average AD of 2.14 ± 0.44 at the genus level and 1.59 ± 0.35 at the species level while JAMS showed 6.03 ± 1.34 at genus level and 10.35 ± 1.28 at species level.

**Table 3A:** Genus

*Genus*
| Pipeline | Source | AD | Sens | FPRA |
| --- | --- | --- | --- | --- |
| Biobakery4 | Amos HiLo | $5.41 \pm 0.17$ | $100.00 \pm 0.00$ | $0.79 \pm 0.10$ |
| jams202212 | Amos HiLo | $8.68 \pm 0.77$ | $100.00 \pm 0.00$ | $0.59 \pm 0.04$ |
| Biobakery4 | Amos Mixed | $3.04 \pm 0.39$ | $100.00 \pm 0.00$ | $3.77 \pm 0.15$ |
| jams202212 | Amos Mixed | $10.33 \pm 1.10$ | $100.00 \pm 3.42$ | $3.83 \pm 0.13$ |
| Biobakery4 | Tourlousse | $2.14 \pm 0.44$ | $94.74 \pm 0.00$ | $6.38 \pm 0.04$ |
| jams202212 | Tourlousse | $6.03 \pm 1.34$ | $100.00 \pm 0.00$ | $0.13 \pm 0.03$ |

**Table 3B:**
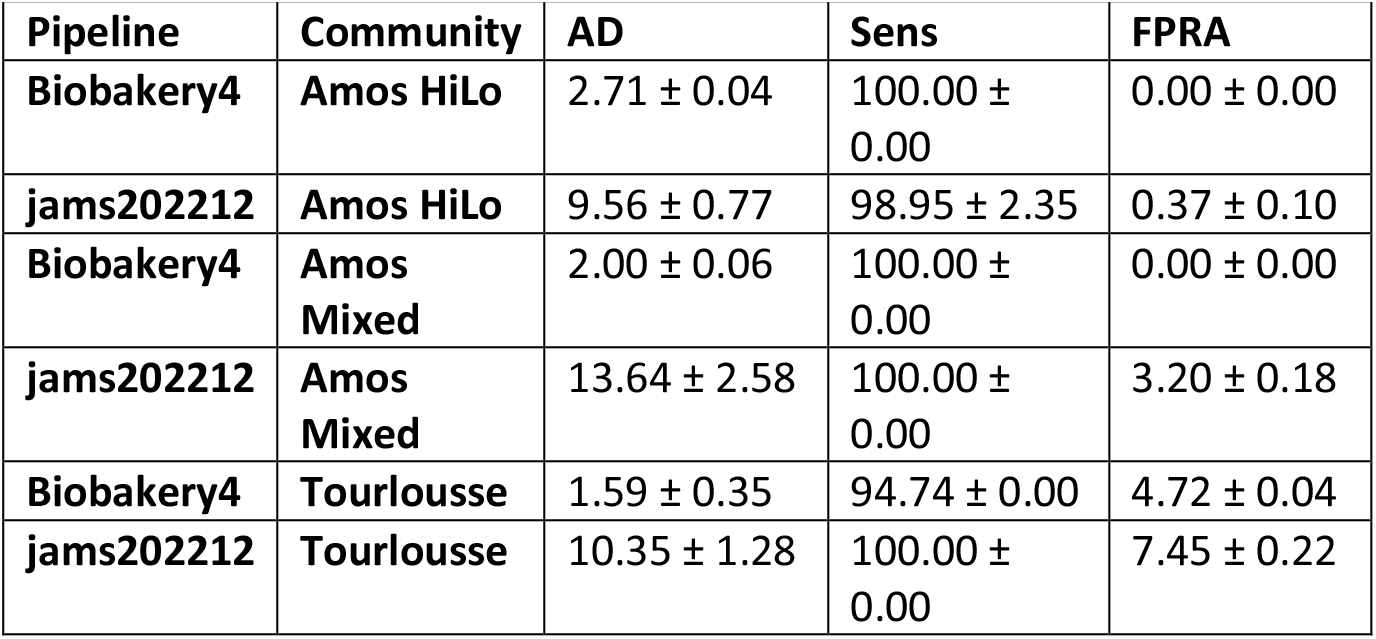
Species

**Figure 8:**
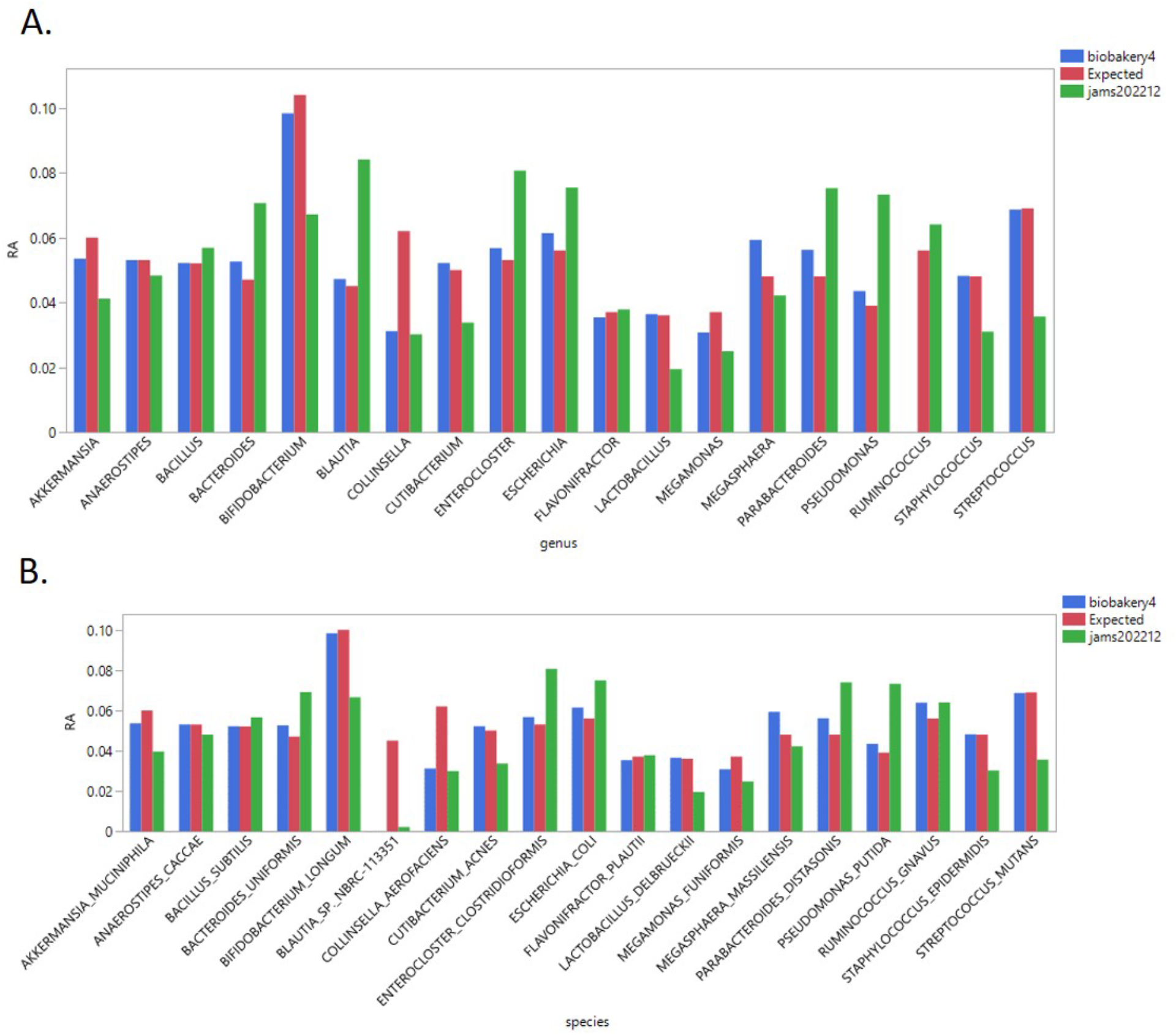
Relative abundance plots of expected and observed for Tourlousse samples at the A) genus level and at the B) species level. Y-axis is the average relative abundance for the expected (red), and the observed relative abundance for Biobakery4 (blue) and JAMS (green). X-axis shows the expected genera and species names.

**Figure 9:**
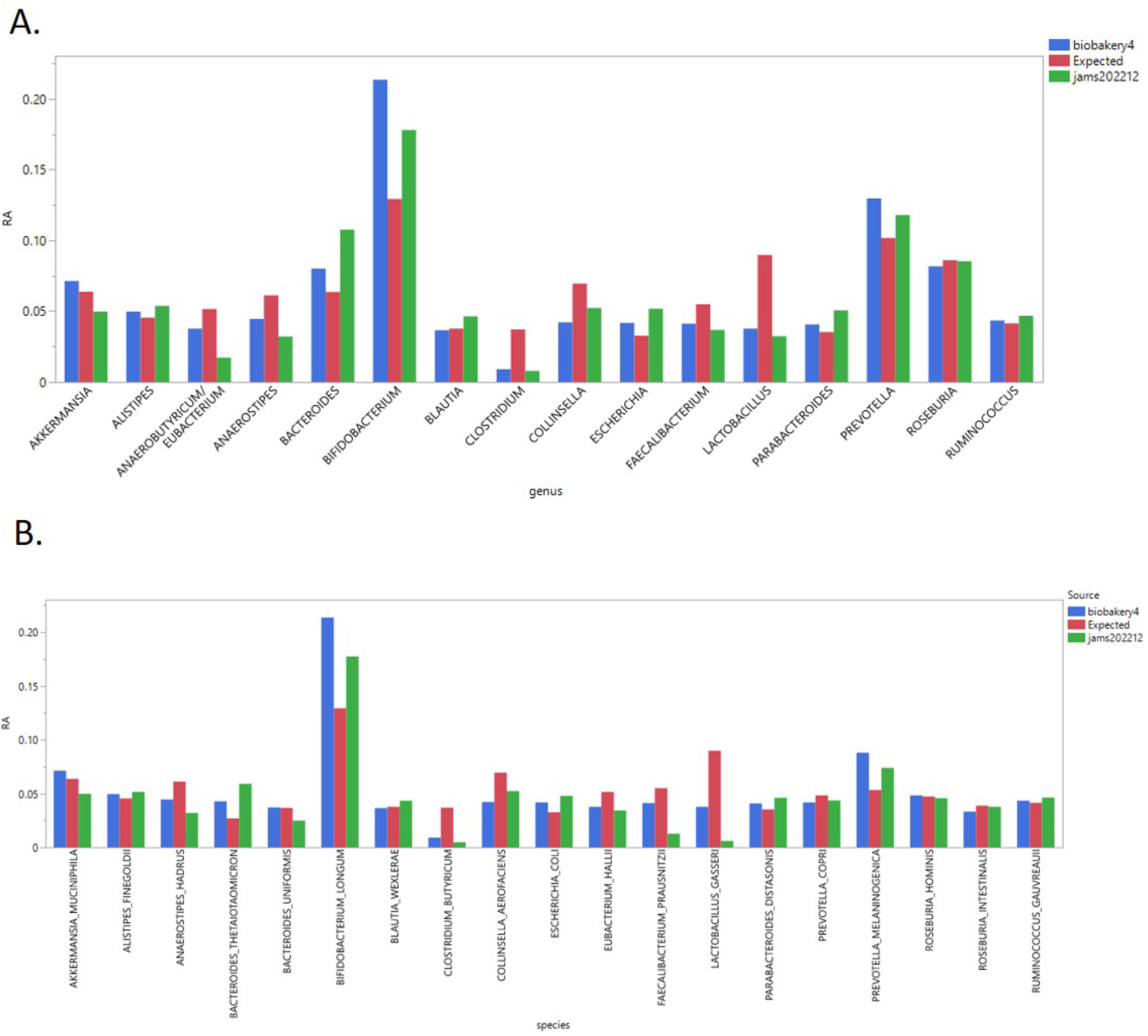
Relative abundance plots of expected and observed for the Amos Mixed samples at the A) genus level and at the B) species level. Y-axis is the average relative abundance for the expected (red), and the observed relative abundance for Biobakery4 (blue) and JAMS (green). X-axis shows the expected genera and species names.

**Figure 10:**
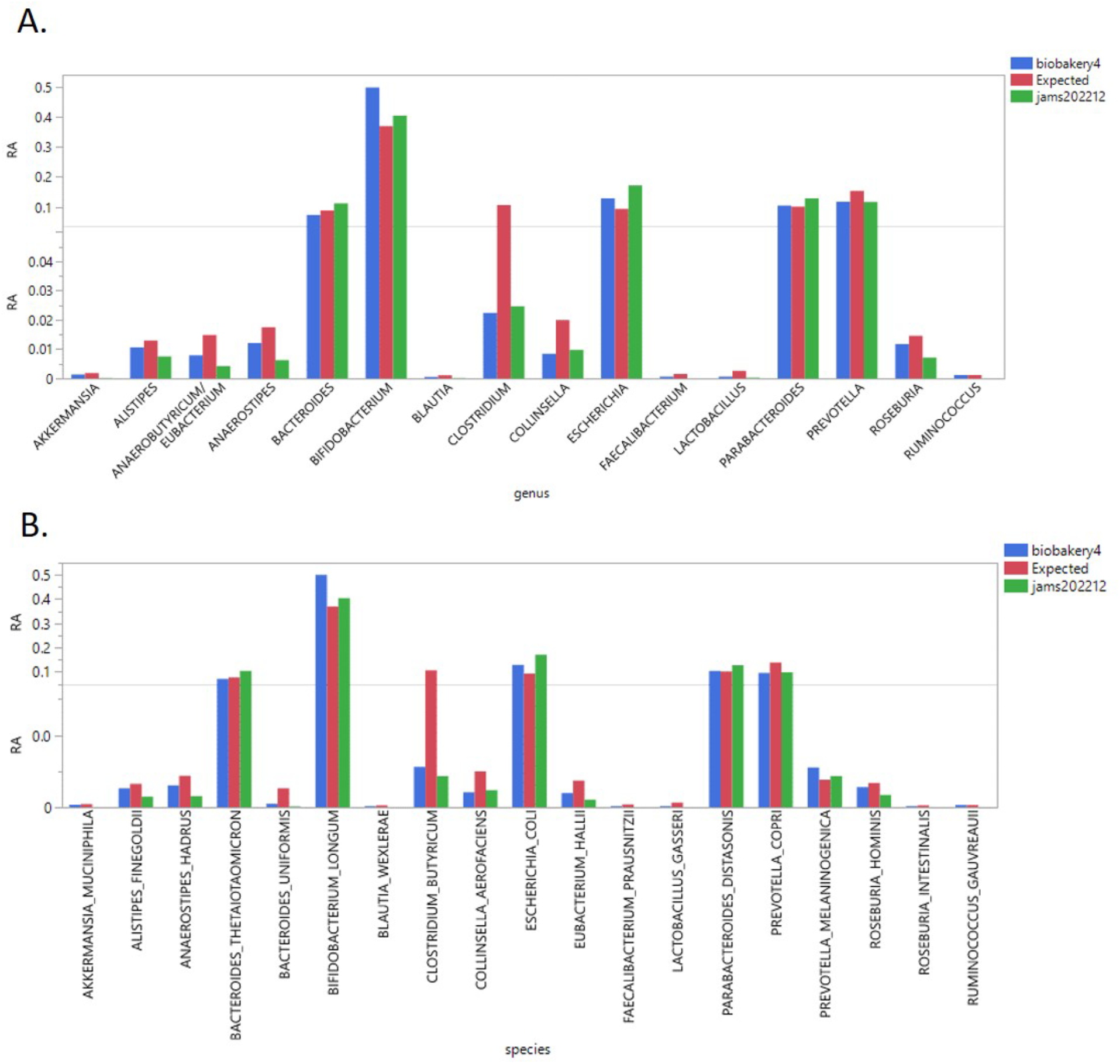
Relative abundance plots of expected and observed for the Amos HiLo samples at the A) genus level and at the B) species level. Y-axis is the average relative abundance for the expected (red), and the observed relative abundance for Biobakery4 (blue) and JAMS (green). X-axis shows the expected genera and species names.

For the Amos Mixed samples, both Biobakery4 and JAMS were able to find all of the expected organisms at the genus and species level. Biobakery4 had 0 FPRA at the species level while JAMS had a total of 3.20 ± 0.18 FPRA at the species level (Table 3B). Biobakery4 showed an average AD of 3.04 ± 0.39 at the genus and 2.00 ± 0.06 at the species while JAMs showed 10.33 ± 1.10 at genus and 13.64 ± 2.58 at species. For the Amos HiLo samples, both Biobakery4 and JAMS were also able to find all 16 of the genera in the mock community samples. Furthermore, they found all the expected species. Note that *Eubacterium halii* has been reclassified as Anaerobutyricum hallii. Since the method of finding genus names is contingent on splitting the genus and species from scientific names, JAMS and Biobakery4 were originally noted as not finding *Eubacterium halii*. This discrepancy was manually corrected. Biobakery4 had 0 FPRA at the species level while JAMS had a total of 0.37 ± 0.10 FPRA at the species level (Table 3B).

Biobakery4 showed an average AD of 5.41 ± 0.17 at the genus and 2.71 ± 0.04 at the species while JAMs showed 8.68 ± 0.77 at genus and 9.56 ± 0.77 at species.

## References

1. Khachatryan, L. et al. Taxonomic classification and abundance estimation using 16S and WGS—A comparison using controlled reference samples. Forensic Sci. Int. Genet. 46, 102257 (2020).

2. Handelsman, J., Rondon, M. R., Brady, S. F., Clardy, J. & Goodman, R. M. Molecular biological access to the chemistry of unknown soil microbes: a new frontier for natural products. Chem. Biol. 5, R245–R249 (1998).

3. Sharon, I., Bercovici, S., Pinter, R. Y. & Shlomi, T. Pathway-Based Functional Analysis of Metagenomes. J. Comput. Biol. 18, 495–505 (2011).

4. Abraham, B. S. et al. Shotgun metagenomic analysis of microbial communities from the Loxahatchee nature preserve in the Florida Everglades. Environ. Microbiome 15, 2 (2020).

5. Frey, B. et al. Shotgun Metagenomics of Deep Forest Soil Layers Show Evidence of Altered Microbial Genetic Potential for Biogeochemical Cycling. Front. Microbiol. 13, (2022).

6. Rosshart, S. P. et al. Laboratory mice born to wild mice have natural microbiota and model human immune responses. Science 365, eaaw4361 (2019).

7. Sato, Y. et al. Taxonomic and functional characterization of the rumen microbiome of Japanese Black cattle revealed by 16S rRNA gene amplicon and metagenome shotgun sequencing. FEMS Microbiol. Ecol. 97, fiab152 (2021).

8. Proctor, L. M. et al. The Integrative Human Microbiome Project. Nature 569, 641–648 (2019).

9. Slatko, B. E., Gardner, A. F. & Ausubel, F. M. Overview of Next Generation Sequencing Technologies. Curr. Protoc. Mol. Biol. 122, e59 (2018).

10. Marcy, Y. et al. Dissecting biological “dark matter” with single-cell genetic analysis of rare and uncultivated TM7 microbes from the human mouth. Proc. Natl. Acad. Sci. U. S. A. 104, 11889–11894 (2007).

11. Rinke, C. et al. Insights into the phylogeny and coding potential of microbial dark matter. Nature 499, 431–437 (2013).

12. Lloyd, K. G., Steen, A. D., Ladau, J., Yin, J. & Crosby, L. Phylogenetically Novel Uncultured Microbial Cells Dominate Earth Microbiomes. mSystems 3, e00055–18 (2018).

13. Ng, P. C. & Kirkness, E. F. Whole Genome Sequencing. in Genetic Variation: Methods and Protocols (eds. Barnes, M. R. & Breen, G.) 215–226 (Humana Press, 2010). doi:10.1007/978-1-60327-367-1_12.

14. Segata, N. et al. Metagenomic microbial community profiling using unique clade-specific marker genes. Nat. Methods 9, 811–814 (2012).

15. Truong, D. T. et al. MetaPhlAn2 for enhanced metagenomic taxonomic profiling. Nat. Methods 12, 902–903 (2015).

16. Ounit, R., Wanamaker, S., Close, T. J. & Lonardi, S. CLARK: fast and accurate classification of metagenomic and genomic sequences using discriminative k-mers. BMC Genomics 16, 236 (2015).

17. Milanese, A. et al. Microbial abundance, activity and population genomic profiling with mOTUs2. Nat. Commun. 10, 1014 (2019).

18. Beghini, F. et al. Integrating taxonomic, functional, and strain-level profiling of diverse microbial communities with bioBakery 3. eLife 10, e65088 (2021).

19. von Meijenfeldt, F. A. B., Arkhipova, K., Cambuy, D. D., Coutinho, F. H. & Dutilh, B. E. Robust taxonomic classification of uncharted microbial sequences and bins with CAT and BAT. Genome Biol. 20, 217 (2019).

20. Weber, N. et al. Nephele: a cloud platform for simplified, standardized and reproducible microbiome data analysis. Bioinformatics 34, 1411–1413 (2018).

21. Meyer, F. et al. Critical Assessment of Metagenome Interpretation: the second round of challenges. Nat. Methods 19, 429–440 (2022).

22. Caporaso, J. G. et al. QIIME allows analysis of high-throughput community sequencing data. Nat. Methods 7, 335–336 (2010).

23. McCulloch, J. A. et al. Intestinal microbiota signatures of clinical response and immune-related adverse events in melanoma patients treated with anti-PD-1. Nat. Med. 28, 545–556 (2022).

24. Davar, D. et al. Fecal microbiota transplant overcomes resistance to anti-PD-1 therapy in melanoma patients. Science 371, 595–602 (2021).

25. Spencer, C. N. et al. Dietary fiber and probiotics influence the gut microbiome and melanoma immunotherapy response. Science 374, 1632–1640 (2021).

26. Stacy, A. et al. Infection trains the host for microbiota-enhanced resistance to pathogens. Cell 184, 615–627.e17 (2021).

27. Hild, B. et al. Neonatal exposure to a wild-derived microbiome protects mice against diet-induced obesity. Nat. Metab. 3, 1042–1057 (2021).

28. Kelsey, C. M. et al. Gut microbiota composition is associated with newborn functional brain connectivity and behavioral temperament. Brain. Behav. Immun. 91, 472–486 (2021).

29. Namasivayam, S. et al. Correlation between Disease Severity and the Intestinal Microbiome in Mycobacterium tuberculosis-Infected Rhesus Macaques. mBio 10, e01018–19 (2019).

30. Fatkhullina, A. R. et al. An Interleukin-23-Interleukin-22 Axis Regulates Intestinal Microbial Homeostasis to Protect from Diet-Induced Atherosclerosis. Immunity 49, 943–957.e9 (2018).

31. Zankari, E. et al. Identification of acquired antimicrobial resistance genes. J. Antimicrob. Chemother. 67, 2640–2644 (2012).

32. Chen, L., Zheng, D., Liu, B., Yang, J. & Jin, Q. VFDB 2016: hierarchical and refined dataset for big data analysis--10 years on. Nucleic Acids Res. 44, D694–697 (2016).

33. Seemann, T. ABRicate. (2023).

34. Bolger, A. M., Lohse, M. & Usadel, B. Trimmomatic: a flexible trimmer for Illumina sequence data. Bioinforma. Oxf. Engl. 30, 2114–2120 (2014).

35. Langmead, B. & Salzberg, S. L. Fast gapped-read alignment with Bowtie 2. Nat. Methods 9, 357–359 (2012).

36. Li, D. et al. MEGAHIT v1.0: A fast and scalable metagenome assembler driven by advanced methodologies and community practices. Methods San Diego Calif 102, 3–11 (2016).

37. Bankevich, A. et al. SPAdes: a new genome assembly algorithm and its applications to single-cell sequencing. J. Comput. Biol. J. Comput. Mol. Cell Biol. 19, 455–477 (2012).

38. Wood, D. E., Lu, J. & Langmead, B. Improved metagenomic analysis with Kraken 2. Genome Biol. 20, 257 (2019).

39. Jones, P. et al. InterProScan 5: genome-scale protein function classification. Bioinforma. Oxf. Engl. 30, 1236–1240 (2014).

40. Li, H. et al. The Sequence Alignment/Map format and SAMtools. Bioinforma. Oxf. Engl. 25, 2078–2079 (2009).

41. Quinlan, A. R. & Hall, I. M. BEDTools: a flexible suite of utilities for comparing genomic features. Bioinforma. Oxf. Engl. 26, 841–842 (2010).

42. Morgan, M., Obenchain, V., Hester, J. & Pagès, H. SummarizedExperiment: SummarizedExperiment container. (2023) doi:10.18129/B9.bioc.SummarizedExperiment.

43. Bates, D. et al. Matrix: Sparse and Dense Matrix Classes and Methods. (2022).

44. Gu, Z., Eils, R. & Schlesner, M. Complex heatmaps reveal patterns and correlations in multidimensional genomic data. Bioinformatics 32, 2847–2849 (2016).

45. Tourlousse, D. M. et al. Characterization and Demonstration of Mock Communities as Control Reagents for Accurate Human Microbiome Community Measurements. Microbiol. Spectr. 10, e01915–21 (2022).

46. Amos, G. C. A. et al. Developing standards for the microbiome field. Microbiome 8, 98 (2020).

47. Babraham Bioinformatics - FastQC A Quality Control tool for High Throughput Sequence Data. https://www.bioinformatics.babraham.ac.uk/projects/fastqc/.

48. Blanco-Miguez, A. et al. Extending and improving metagenomic taxonomic profiling with uncharacterized species with MetaPhlAn 4. 2022.08.22.504593 Preprint at https://doi.org/10.1101/2022.08.22.504593 (2022).

49. Gloor, G. B., Macklaim, J. M., Pawlowsky-Glahn, V. & Egozcue, J. J. Microbiome Datasets Are Compositional: And This Is Not Optional. Front. Microbiol. 8, (2017).

50. Lubbe, S., Filzmoser, P. & Templ, M. Comparison of zero replacement strategies for compositional data with large numbers of zeros. Chemom. Intell. Lab. Syst. 210, 104248 (2021).

